# Longitudinal allometry of sulcal morphology in health and schizophrenia

**DOI:** 10.1101/2021.03.17.435797

**Authors:** Joost Janssen, Clara Alloza, Covadonga M. Díaz-Caneja, Javier Santonja, Laura Pina-Camacho, Pedro M. Gordaliza, Alberto Fernández-Pena, Noemi González Lois, Elizabeth E.L. Buimer, Neeltje E.M. van Haren, Wiepke Cahn, Eduardo Vieta, Josefina Castro-Fornieles, Miquel Bernardo, Mara Parellada, Celso Arango, René S. Kahn, Hilleke E. Hulshoff Pol, Hugo G. Schnack

## Abstract

Scaling between subcomponents of cortical folding and total brain volume (TBV) in healthy individuals (HI) is allometric, i.e. non-linear. It is unclear whether this is also true in individuals with schizophrenia (SZ) or first-episode psychosis (FEP). The current study first confirmed normative allometric scaling norms in HI using discovery and replication samples. Cross-sectional and longitudinal diagnostic differences in folding subcomponents were then assessed using an allometric analytic framework.

Structural imaging from a longitudinal (sample 1: HI and SZ, n_HI Baseline_ = 298, n_SZ Baseline_ = 169, n_HI Follow-up_ = 293, n_SZ Follow-up_ = 168, a total of 1087 images, all individuals ≥ 2 images, age 16-69 years) and a cross-sectional sample (sample 2: n_HI_ = 61 and n_FEP_ = 89, age 10-30 years) is leveraged to calculate global folding and its nested subcomponents: sulcation index (SI, total sulcal/cortical hull area) and determinants of sulcal area; sulcal length and sulcal depth.

Scaling of the SI, sulcal area, and sulcal length with TBV in SZ and FEP was allometric and did not differ from HI. Longitudinal age trajectories demonstrated steeper loss of SI and sulcal area through adulthood in SZ. Longitudinal allometric analysis revealed that both annual change in SI and sulcal area was significantly stronger related to change in TBV in SZ compared to HI.

Our results detail the first evidence of the disproportionate contribution of changes in SI and sulcal area to TBV changes in SZ. Longitudinal allometric analysis of sulcal morphology provides deeper insight into lifespan trajectories of cortical folding in SZ.

## Introduction

The degree of folding of the human cortex is an important marker of early-life neurodevelopmental processes (Llinares-Benadero and Borrell, 2019). Early-life neurodevelopmental processes are thought to be aberrant in first-episode psychosis (FEP) and schizophrenia (SZ). This notion is corroborated by consistently reported average reductions in the degree of folding and sulcation of FEP and SZ relative to healthy individuals (HI) (Penttilä et al., 2009; Palaniyappan et al., 2011; Janssen et al., 2014; Lavoie et al., 2014; Nanda et al., 2014; Nesvåg et al., 2014; Cachia et al., 2015; Gay et al., 2017; MacKinley et al., 2019; Dazzan et al., 2020). However, the degree of folding is not static throughout life in HI and reduces postnatally, during adolescence, and beyond (Alemán-Gómez et al., 2013; Mutlu et al., 2013; Li et al., 2014; Cao et al., 2017).

Longitudinal studies are necessary to establish whether SZ is associated with abnormalities in folding reductions and whether the pattern of SZ-related abnormalities may be dynamic over time. In addition, longitudinal studies can clarify whether SZ-related folding abnormalities can also occur at later stages of the illness during late adulthood and old age (Mirakhur et al., 2009; Raznahan et al., 2011; Li et al., 2014).

The in vivo degree of folding can be estimated using the gyrification index or the sulcation index (SI) (Shimony et al., 2016). These metrics are composite scores that combine two components: hull surface area (exposed cortex) and outer cortex in the case of the gyrification index and sulcal surface area (non-exposed cortex) in the case of SI. Sulcal surface area can in turn can be decomposed into sulcal depth and length allowing for a complete decomposition of the SI into folding subcomponents (see Figure 1) (Mangin et al., 2004; Alemán-Gómez et al., 2013). Assessing the SI and its subcomponents in a longitudinal lifespan study of HI and individuals with SZ allows for a fine-grained assessment of change in the degree of folding across the lifespan and whether SZ contributes distinctively to the folding subcomponents.

**Figure 1.**
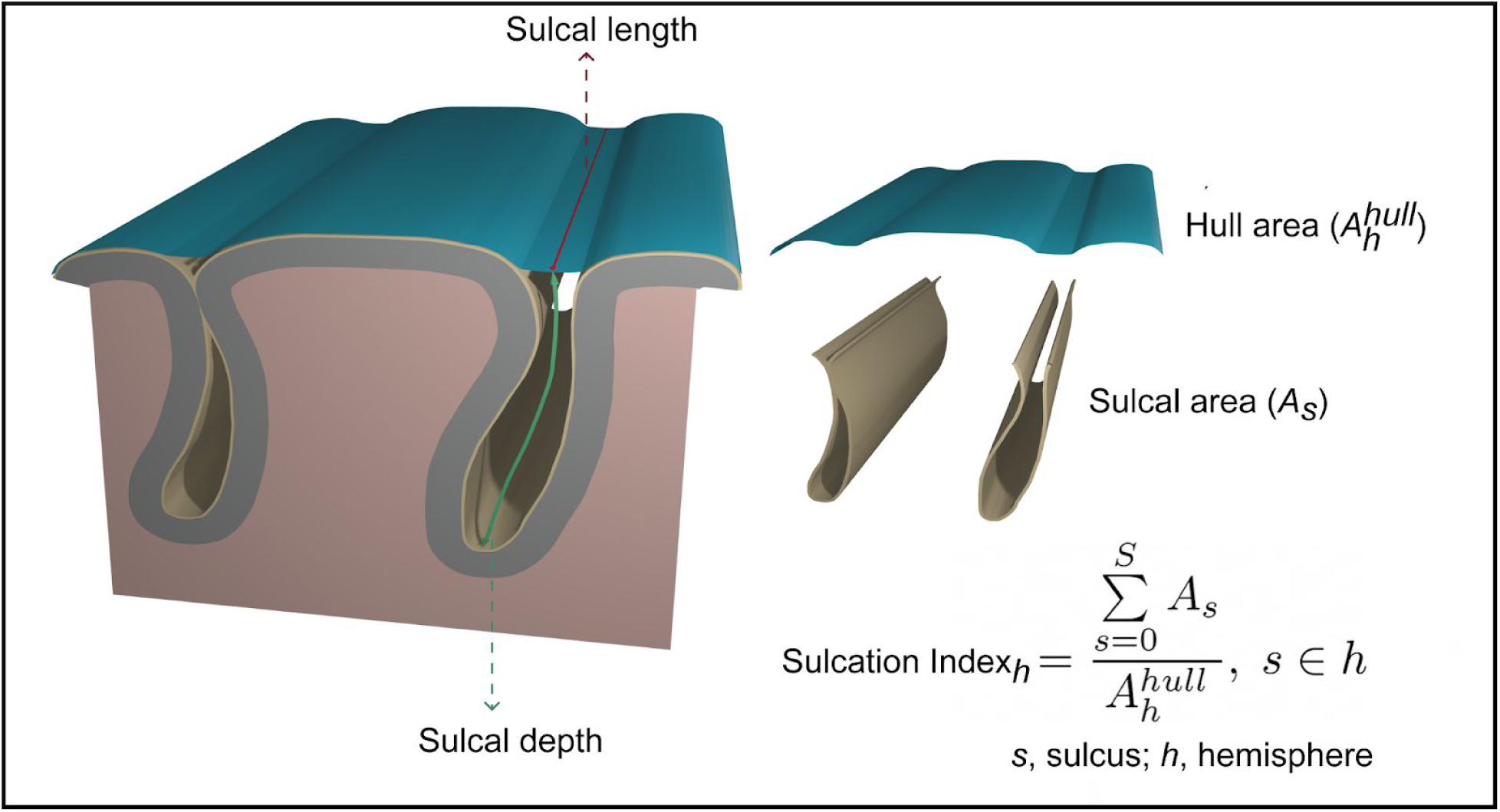
Graphical representation of SI and its components. Total sulcal surface is defined as the sum of the area of all sulcal median meshes. Hull area is derived by calculating the area of the brain mesh defined via a morphologic closing of the brain mask which excludes sulcal areas during definition of the exposed cortical surface area. Total sulcal length is defined as the sum of the length of all median sulcal meshes. Geodesic sulcal depth is the shortest distance along the surface of the brain from the point to where the brain surface makes contact with the hull surface.

Reduced total brain volume (TBV) is a robust neurobiological feature of psychotic disorders (Haijma et al., 2013). Since smaller brain sizes are associated with a lower degree of folding and early-life folding growth positively correlates with brain volume growth, the question arises whether reduced folding in psychotic disorders may be merely due to a global decrease in brain size (Toro et al., 2008). Accounting for brain size in comparisons of folding metrics between individuals with psychotic disorders and HI is usually done using traditional methods (e.g. including TBV as a nuisance variable). However, these approaches assume an identical linear association between folding and TBV across groups. However, the relationship between brain metrics may be allometric, i.e. non-linear (Im et al., 2008; Toro et al., 2008; Barnes et al., 2010; Reardon et al., 2016; de Jong et al., 2017; Mankiw et al., 2017; Jäncke et al., 2019). Using a HI sample in an allometric framework allows for providing normative scaling norms between folding subcomponents and TBV. These norms are necessary for determining whether the effect of SZ on subcomponents of folding is due to scaling differences or whether folding abnormalities are present independently from group differences in TBV.

Allometric scaling laws predict that the static relationship between brain volume and areal measures (sulcal and hull area) in a log-log framework should be ⅔ while the relationship with linear measures (sulcal depth and length) should be ⅓ (Toro et al., 2008; Fish et al., 2017). However, the only study so far to assess scaling of brain volume with sulcal folding metrics in neurotypical children show that larger brains have a higher SI due to a disproportionate increase in sulcal length but not depth (Fish et al., 2017). While interesting, it is unclear whether the observed allometric scaling also affects longitudinal trajectories of folding metrics and modulates potential differences in trajectories between SZ and HI.

In this study we first set out to replicate the previously reported normative scaling norms between TBV and the folding metrics in two independent samples of HI. Thereafter, we assess separately in each sample whether SZ or FEP groups differ from the HI group in allometric scaling. We then compare longitudinal age trajectories of folding metrics between SZ and HI. Finally we apply we longitudinal allometric scaling to examine how change in folding subcomponents relate to change in TBV and whether this scaling differs between SZ and health.

## Materials and Methods

Our study uses two independent samples (sample 1, sample 2). Detailed clinical, demographic and imaging measures for both samples are summarized in Table 1 and 2.

**Table 1.**
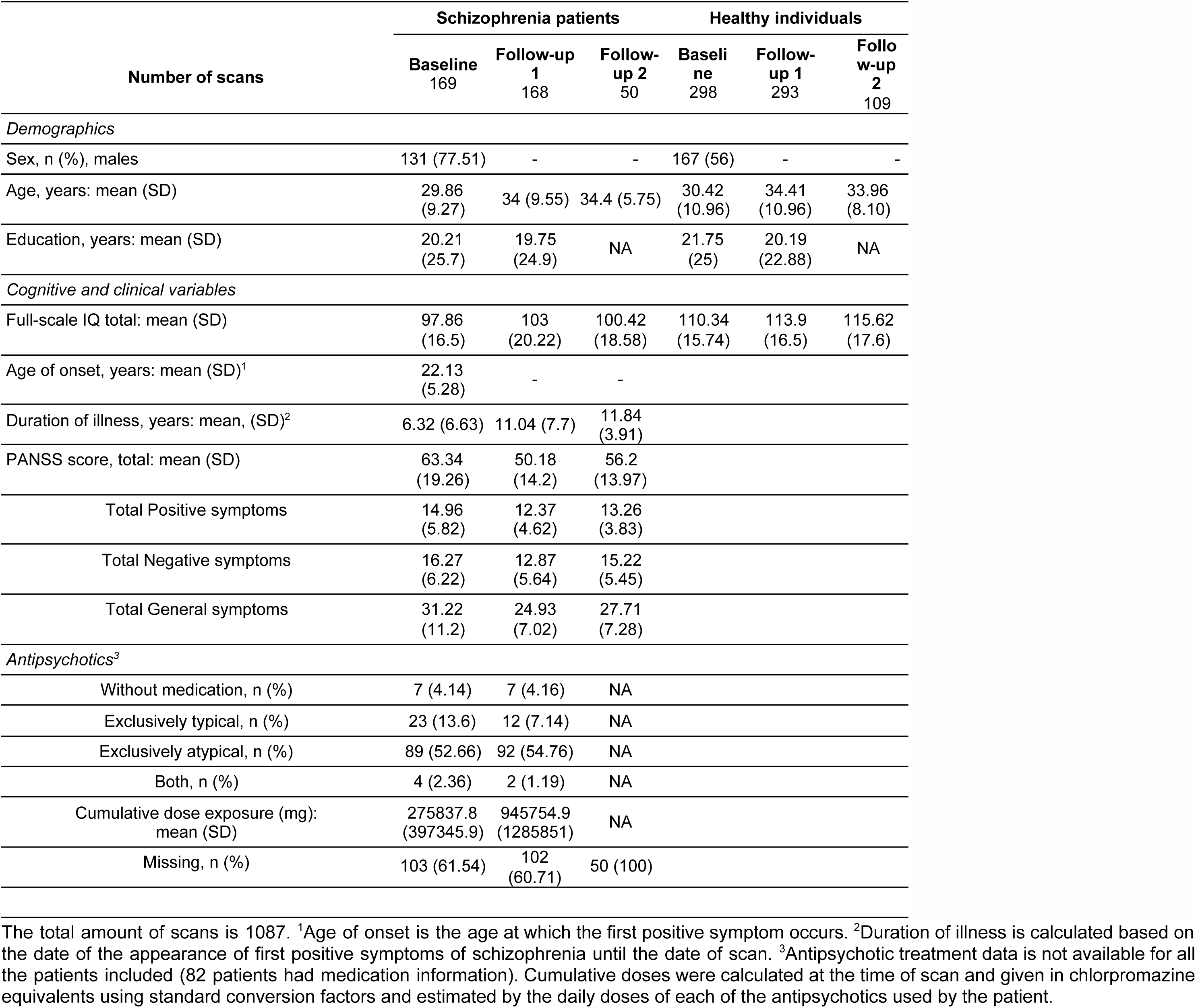
Demographic, cognitive and clinical characteristics of individuals with schizophrenia and healthy individuals for sample 1.

**Table 2.**
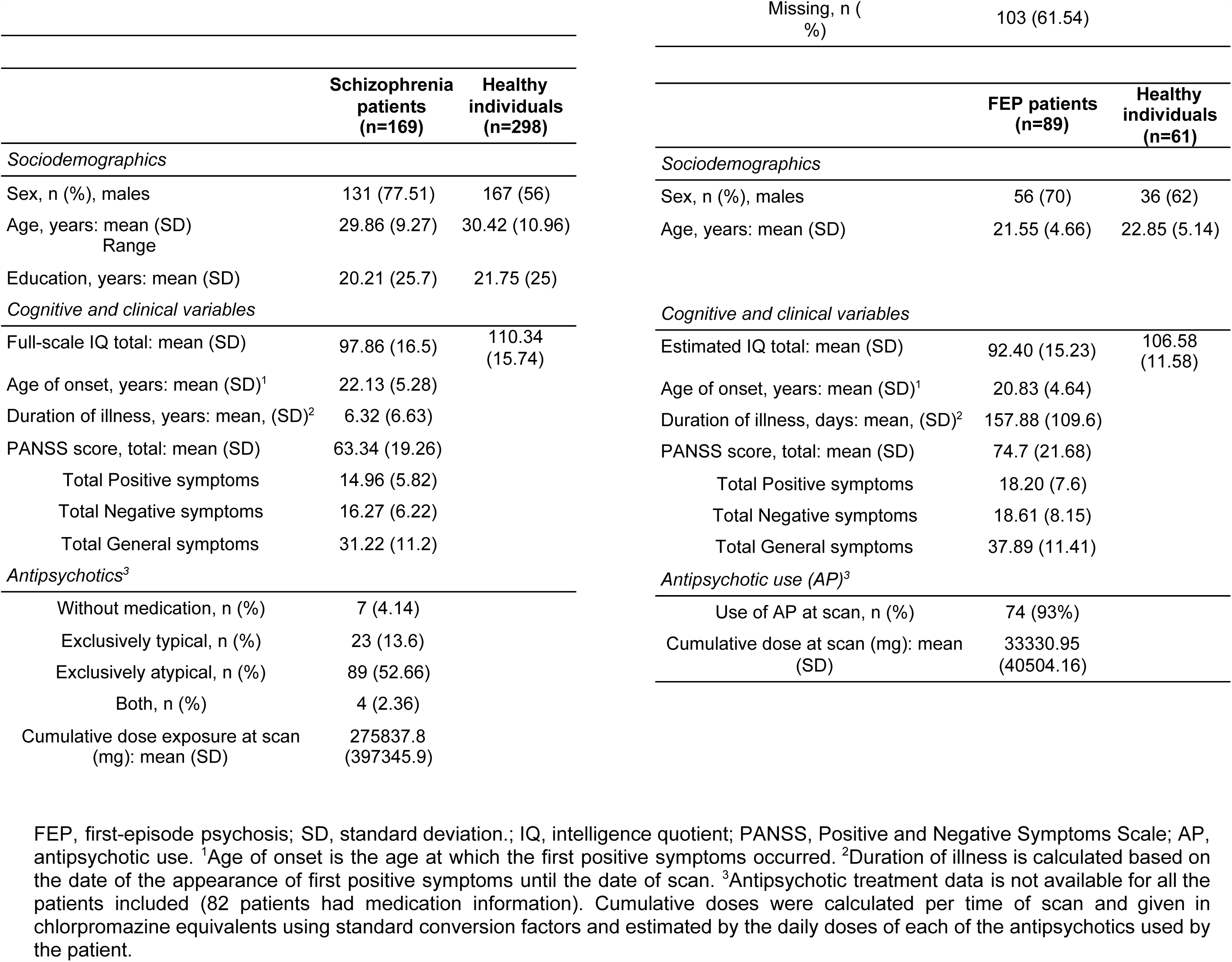
Demographic, cognitive and clinical variables of individuals with schizophrenia or first-episode psychosis and healthy individuals of samples 1 (at baseline) and 2.

### Sample 1: schizophrenia and healthy individuals (longitudinal)

From a large longitudinal sample of individuals with schizophrenia and HI aged 16-70 years at baseline we included individuals who had T1-weighted magnetic resonance imaging (MRI) scan acquisitions at two or three time points. Detailed information regarding diagnostic criteria and clinical and cognitive assessments of the Utrecht Schizophrenia project and the Genetic Risk and Outcome of Psychosis (GROUP) consortium are described in (Hulshoff Pol et al., 2001; Korver et al., 2012; Kubota et al., 2015). Detailed information on the current subsample can be found in (Janssen et al., 2020). Details on the MRI quality assessment (both FreeSurfer and BrainVISA) and time between scans can be found in the Supplemental text and SFigures 1 and 2. The final dataset consisted of 1087 scans from 298 HI and 168 patients. Demographic, cognitive and clinical information of the participants at all time points is summarized in Table 1 and information at baseline is summarized in Table 2 (left column). The study was approved by the institutional review board and was conducted according to the provisions of the World Medical Association Declaration of Helsinki. Written informed consent was obtained from all participants and also from parents/legal guardians for children under 18 years of age.

### Sample 2: first-episode psychosis and healthy individuals (cross-sectional)

The sample is a subset of participants with available imaging data from a larger multi-site, naturalistic study including participants aged 7-35 years with a first episode of psychosis, the PEPs Project. The main objective of the PEPs project was to identify the several gene-by-environment interactions involved in the risk of psychosis in a naturalistic cohort of first-episode psychosis to develop a predictive model of psychosis. Most of the centers were integrated in the well-recognized Spanish network of research in Mental Health named “Centro de Investigación Biomédica en Red de Salud Mental (CIBERSAM)”. Detailed information about the study design, recruitment procedures, in- and exclusion criteria, clinical and cognitive assessments, are provided elsewhere (Bernardo et al., 2013; Pina-Camacho et al., 2016). For the current study only participants scanned at a single site (Barcelona) were included (n=182) of which a further 32 participants were excluded based on image quality. Detailed demographic, cognitive and clinical information for the final sample is summarized in Table 2 (right column). Details on the MRI quality assessment can be found in the Supplemental text. The study was approved by the institutional review board and was conducted according to the provisions of the World Medical Association Declaration of Helsinki. Written informed consent was obtained from all participants and from parents/legal guardians for children under 18 years of age.

## MRI acquisition

### Image acquisition

For sample 1, two scanners (same vendor, field strength and acquisition protocol) were used. Participants were scanned twice on either a Philips Intera or Achieva 1.5 T and a T1-weighted, 3-dimensional, fast-field echo scan with 160-180 1.2 mm contiguous coronal slices (echo time [TE], 4.6 ms; repetition time [TR], 30 ms; flip angle, 30°; field of view [FOV], 256 mm; in-plane voxel size, 1×1 mm^2^) was acquired. All included participants had their baseline and follow-up scan on the same scanner. For sample 2, participants were scanned on a Siemens Trio TIM 3 T and a T1-weighted, 3-dimensional, gradient echo scan with 240 1 mm contiguous sagittal slices (echo time [TE], 2.98 ms; repetition time [TR], 2300 ms; flip angle, 9°; field of view [FOV], 256 mm; in-plane voxel size, 1×1 mm^2^) was acquired.

### Image analysis

TBV, SI, total sulcal and hull area, global average sulcal depth, and total sulcal length were assessed (see Figure 1). All images were analyzed using the FreeSurfer analysis suite (v5.1) with default settings to provide total gray and white matter volumes which were summed to calculate TBV (Dale et al., 1999; Fischl et al., 1999; Fischl, 2012). For all images, sulcal segmentation and identification was performed with BrainVISA software (v4.5) using the Morphologist Toolbox and Mindboggle software using default settings (Mangin et al., 2004, 2010; Klein et al., 2017). After importing Freesurfer’s ribbon image into BrainVISA each sulcus is segmented with the cortical sulci corresponding to the crevasse bottoms of the “landscape,” the altitude of which is defined by image intensity. A spherical hull surface was generated from a smooth envelope that wrapped around the hemisphere but did not encroach into the sulci, a morphological isotropic closing of 6 mm was applied to ensure boundary smoothness. The median sulcal surface spans the entire space contained in a sulcus, from the fundus to its intersection with the hull. For each fold, sulcal area is defined as the total surface area of the medial sulcal surface, sulcal length is measured on the hull and is defined as the distance of the intersection between the median sulcal surface and the hull. All sulcal nodes belonging to a hemisphere were relabeled to one label (see SFigure 1) and then BrainVISA software was used to calculate hemispheric sulcal area and length. FreeSurfer output was imported into Mindboggle software for calculating geodesic sulcal fundi depth. All metrics were measured in the native space of the participant’s images and left and right hemisphere values were either summed or averaged. FreeSurfer, BrainVISA and Mindboggle derived measurements have been validated via histological and manual measurements and have shown good test–retest reliability (Rosas et al., 2002; Kuperberg et al., 2003; Pizzagalli et al., 2020).

## Statistics

All analyses were performed in R (https://cran.rstudio.com/).

### Allometric analyses at baseline: samples 1 and 2

#### Normative scaling coefficients from healthy individuals

We first regressed out the effects of age (age, age^2^, age^3^), sex, and scanner (only for sample 1), by computing at each timepoint separately a linear regression model with each metric *m* as dependent variable and the covariates as predictors. To control for collinearity among the polynomials of age we made use of the poly()-function in R which created three uncorrelated variables to model age effects (so-called orthogonal polynomials) (34) for metric *m* (Chambers and Hastie, 1992):

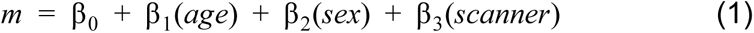

We then saved the residuals and added the mean back in for interpretability.

We converted the residualized metrics from HI (samples 1 and 2 separately) at baseline to log values and applied orthogonal regression to calculate normative baseline allometric scaling coefficients of log transformed values of TBV with SI, sulcal and hull area, sulcal depth, and length. In our scenario, both dependent and independent variables have similar measurement error. Therefore, in our case non-orthogonal regression may lead to biased regression estimates as only the measurement error of the independent variable is taken into account. In orthogonal regression the measurement error of both dependent and independent variables are accounted for, leading to higher accuracy of regression estimates (Madansky, 1959). This method allowed us to assess whether the normative baseline allometric scaling relationships between the log transformed TBV and folding subcomponents were similar in the discovery sample (sample 1) and the replication sample (sample 2) and those reported by Fish et al. (2017) and whether they were similar compared to the expected scaling coefficients. The *t* for the difference between the observed and expected coefficients was calculated as follows:

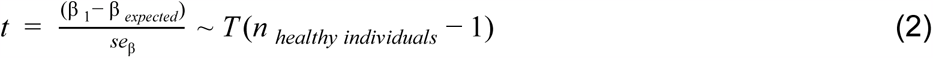

where β_1_ is the scaling coefficient for the HI group, β*_expected_* is the scaling coefficient as expected by scaling laws, *se*_β_ is the standard error of the slope of the HI group given by bootstrapping and *n* is the number of participants. For each folding metric, scaling coefficients below and above the expected values were considered to indicate hypo- and hyperallometry, respectively. Coefficients that were statistically indistinguishable from the null indicated isometric scaling. The *mcr* R package was used for orthogonal regression.

#### Comparison of allometric scaling coefficients between healthy individuals and SZ (sample 1) and FEP (sample 2)

We converted the residualized metrics from SZ and FEP patients (sample 1 and 2 separately) to log values and applied orthogonal regression to compare baseline allometric scaling coefficients between HI and SZ (sample 1) and FEP (sample 2) separately. Significance of the slope was determined through 95% bias-corrected and accelerated bootstrap (BCa) confidence intervals generated using bootstrapping (n=1000) (Efron and Tibshirani, 1993). If the slopes of the SZ and healthy participant groups were both significant we calculated a *t* for the difference between the slopes of the patient and healthy participant groups, as follows:

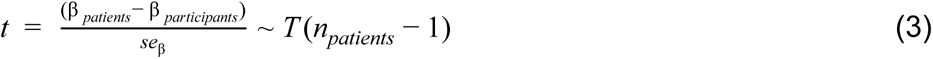

where β*_healthy individuals_* is the scaling coefficient for the HI group, β*_patients_* is the coefficient for the SZ group, *se*_β_ is the standard error of the slope of the SZ group given by the bootstrap and *n_patients_* is the number of patients.

In order to directly compare the results of our allometric approach to traditional methods of controlling for effects of TBV we ran linear models comparing groups with expressing the residualized fold metrics as a fraction of TBV, i.e. normalization:

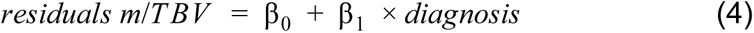

and with adding TBV as a covariate,

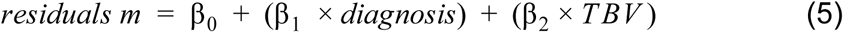

where *m* is the metric.

### Longitudinal age trajectories in SZ and healthy individuals (sample 1)

We used generalized additive mixed models (GAMMs) to investigate the longitudinal trajectories of SI, sulcal and hull area, sulcal depth, and length over the age range (Wood, SN, 2006). We specified cubic regression splines with shrinkage (a type of penalized splines) as smooth terms and set k = 4 as the number of knots of the spline. GAMM models were implemented to examine diagnosis (i.e., SZ vs HI), the age×diagnosis interaction, and TBV, while controlling for sex and scanner (as fixed effects) and the random effect of the individual. Including a dummy variable to account for scanner type as a random effect in the models did not change the results. In order to account for the potential effect of a non-linear relationship between cortical folding metrics and TBV on age trajectories we reran the GAMM models but now transformed TBV and the folding metrics to log values. In these models we also tested whether the log(TBV)×diagnosis interaction was significant. If this interaction was not significant it was omitted from the model. To better understand the age×diagnosis interaction, GAMM estimates for age were also implemented and visualized for patients and HI separately. We used the *mgcv* R package for modeling of the GAMMs. Finally, we plotted a difference fit (predicted SZ fit - predicted HI fit) with its 95% confidence interval against age. The difference fit allows for assessment of where along the age range the difference between SZ and HI becomes significantly different from zero.

### Longitudinal allometric analysis: the relationship between annual change in fold metrics and annual change in brain volume in SZ and healthy individuals (sample 1)

In these analyses we assessed whether yearly changes in folding metrics scaled with yearly changes in TBV and whether this scaling differed between the SZ and HI groups. For this, we used the residuals from baseline and the nearest available follow-up measurement from each subject to determine the symmetrized annual percent change (APC) for each subject and metric *m*:

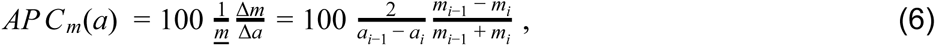

where Δ*m* represents change in metric *m*, obtained at two different time points, *i* and *i* − 1, *i* ∈ [1, 2, 3] (up to three measures per subject in this study) and Δ*a* the time lapse between them in years; *a* is the subject’s age in years at a specific time point. Finally, 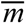 represents the average of the metric’s value at the two time points. We then calculated longitudinal allometric scaling coefficients of the APC of TBV versus the APCs of the folding metrics using orthogonal regression in HI and individuals with SZ. Significance of the slope was determined through 95% bias-corrected and accelerated bootstrap (BCa) confidence intervals generated using bootstrapping (n=1000) (Efron and Tibshirani, 1993). If the slopes of the SZ and HI groups were both significant we calculated a *t* for the difference between the slopes of the patient and healthy participant group using the previously described formula (3).

In all analyses we controlled for multiple comparisons using Bonferroni correction.

## Results

### Allometric analyses at baseline: sample 1 and 2

#### Normative scaling coefficients from healthy individuals

As can be seen in Figure 2 the cross-sectional allometric findings replicated across the two independent samples and replicated the normative findings reported by Fish et al. (2017). In the healthy participant groups of sample 1 and 2, a larger SI was associated with a larger TBV. In both samples there was hyperallometric scaling of SI, sulcal area and length and isometric scaling of hull area. Sulcal depth showed either hypoallometric scaling (sample 1) or isometric scaling (sample 2).

**Figure 2.**
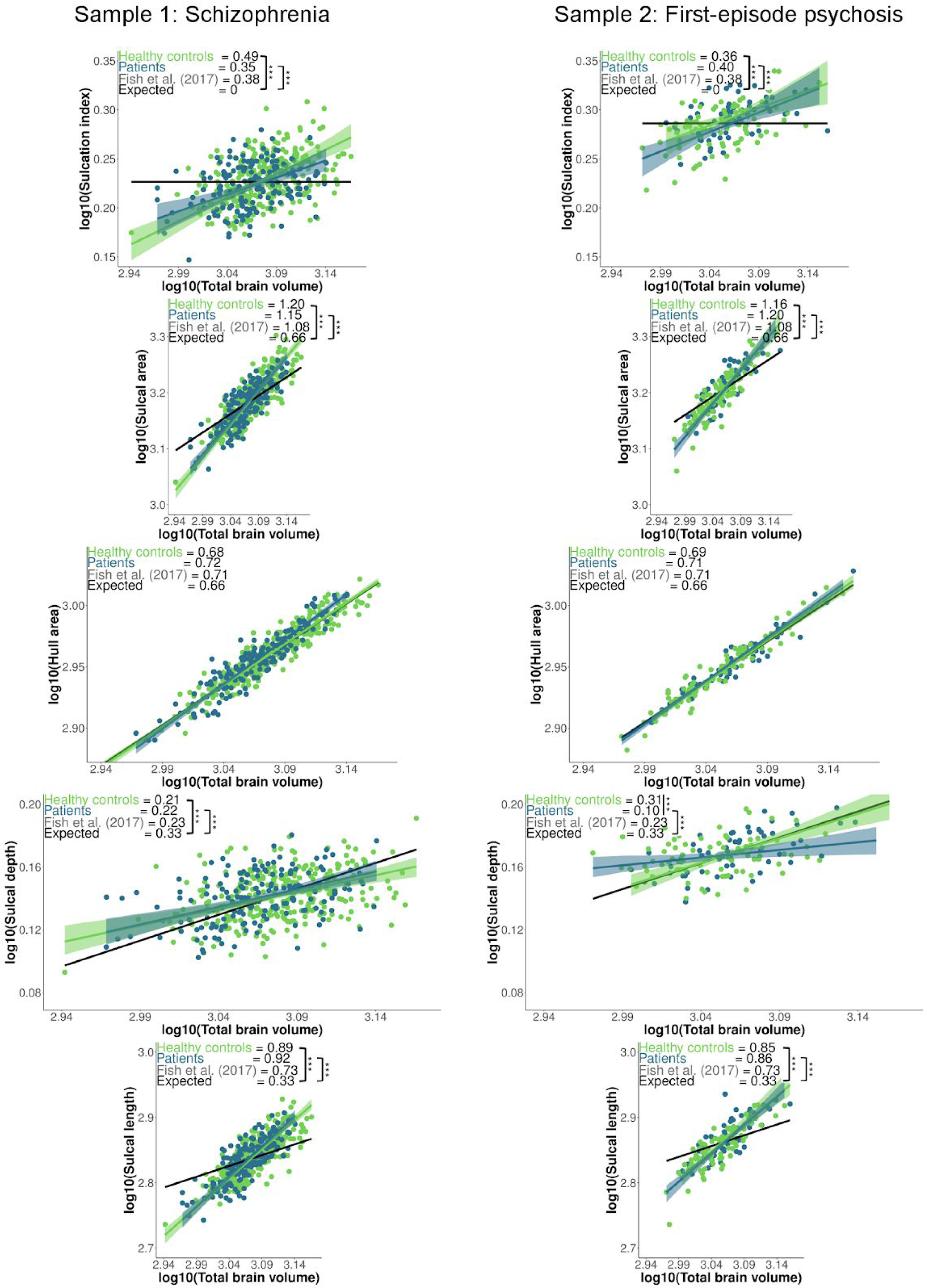
Orthogonal regression fits ± 95% confidence interval and the scaling coefficient for observed log–log allometric scaling of residualized folding metrics relative to residualized total brain volume in the healthy individuals and patient groups at baseline (samples 1 (left) and 2 (right) separately). “Expected” values (black line) represent the expected scaling (from scaling laws) coefficient. ***=p<0.001 denotes significance after Bonferroni correction for the difference between (1) the scaling of the healthy individuals groups and the expected scaling and (2) the healthy individuals groups and patient groups. Values were residualized for scanner (only sample 1), age and sex.

#### Comparison of baseline allometric scaling coefficients between healthy individuals and SZ (sample 1) and FEP (sample 2)

Allometric analyses indicated that in both the SZ and FEP patient groups the SI, sulcal and hull area, and sulcal length were in proportion with normative decreases in TBV (see Figure 2). Allometric analysis clearly demonstrated that SI, sulcal area, and sulcal length scaled non-linearly with TBV in HI, SZ and FEP groups. Traditional methods of “controlling for” TBV differences, which assume linear scaling between the fold metrics and TBV, were not consistent with the results of our allometric analysis (see Table 3). There were no significant associations between the residualized brain metrics, CPZ equivalents, and illness duration. In the group of SZ patients, estimated IQ explained about 9% of the variance of the brain metrics, except for sulcal depth (see SFigure 3).

**Table 3.** Comparing different methods of controlling for brain size in analysis of diagnostic differences at baseline.

|  | Schizophrenia<br>vs control | First-episode psychosis<br>vs control |
| --- | --- | --- |
| Sulcation Index |  |  |
| Normalization | ↑ | — |
| Covarying | — | ↓ |
| Allometry | — | — |
| Sulcal area |  |  |
| Normalization | — | — |
| Covarying | — | — |
| Allometry | — | — |
| Hull area |  |  |
| Normalization | ↑ | ↑ |
| Covarying | ↑ | — |
| Allometry | — | — |
| Sulcal depth |  |  |
| Normalization | ↑ | — |
| Covarying | — | — |
| Allometry | — | ↓ |
| Sulcal length |  |  |
| Normalization | ↑ | — |
| Covarying | — | — |
| Allometry | — | — |

### Longitudinal age trajectories in SZ and healthy individuals (sample 1)

There was no main effect of diagnosis for any of the metrics (see Table 4). The healthy control and SZ group showed loss of TBV, sulcal area, and sulcal depth and length across the lifespan (see Figure 3A). After Bonferroni correction the age×diagnosis interaction was significant for SI and sulcal area (see Table 4). Trajectory differences in SI were larger than zero from 32 years onwards approximately (approximate patient group deficit at 32 years and at the maximum age: −0.02 and −0.08, respectively) and for sulcal area from 44 years onwards, approximately (approximate patient group deficit at 44 years and at the maximum age: 18 and 35cm^2^, respectively), see Figure 3B.

**Table 4.**
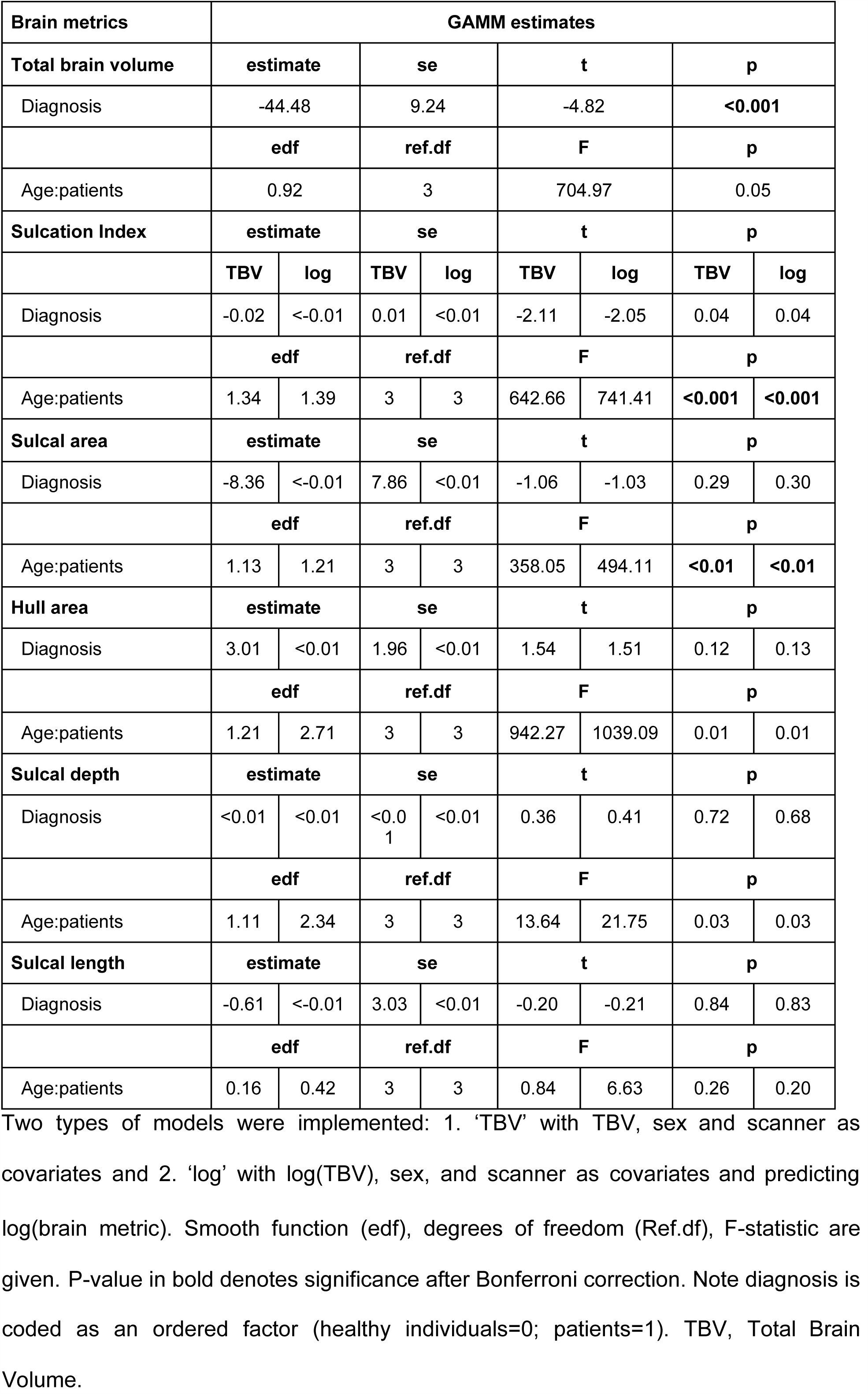
Generalized Additive Mixed Model (GAMM) estimates for diagnosis and age*diagnosis for each brain metric in sample 1.

**Figure 3.**
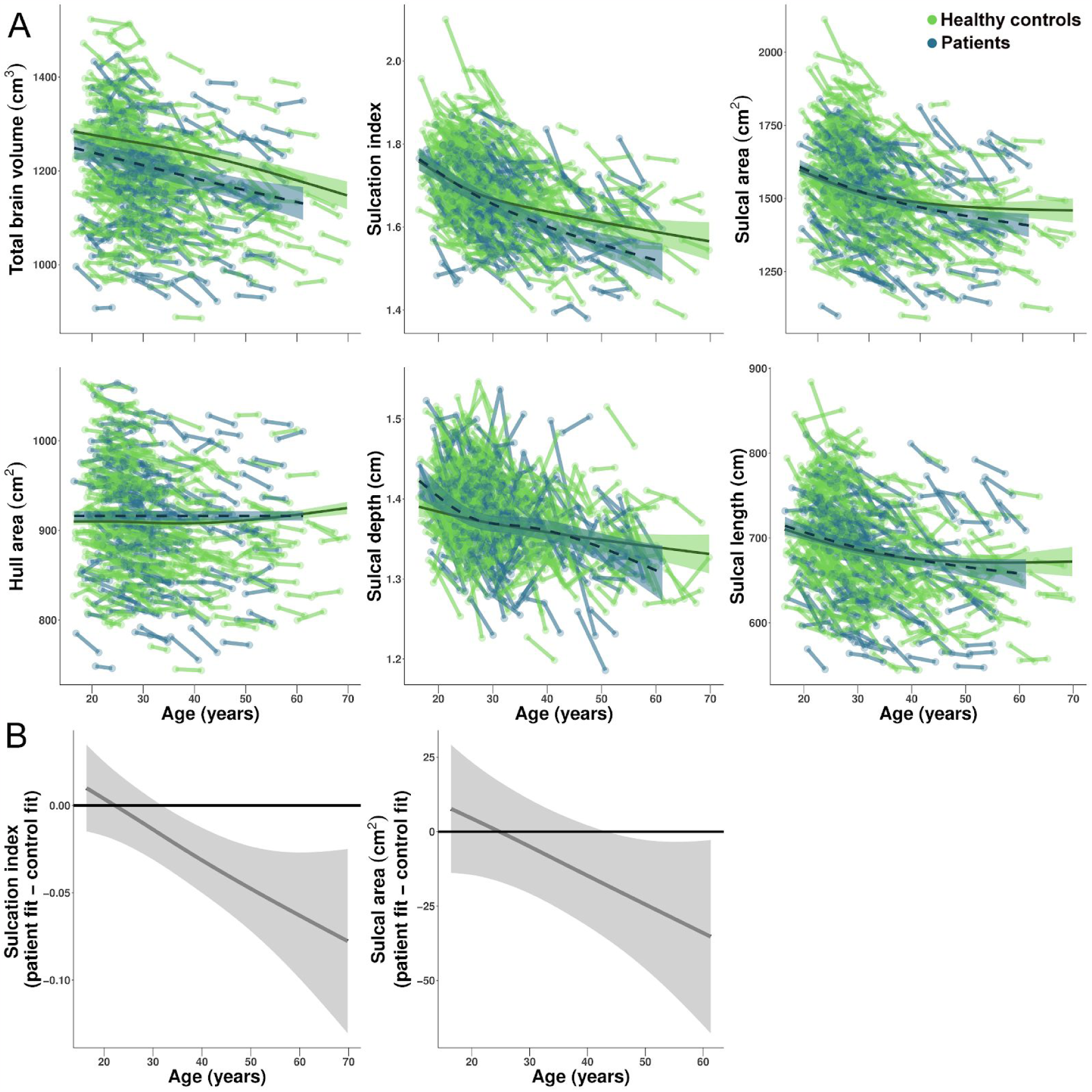
**A**. Age trajectories of total brain volume and the five folding metrics in healthy individuals and individuals with schizophrenia from sample 1. Generalized Additive Mixed Model (GAMM) fits ± 95% confidence intervals are shown over the data points for all participants in the corresponding color, with data at each of the time points connected for each participant. Sex, total brain volume, and scanner were included in the models as covariates (except for the model for total brain volume) and the individual was modeled as a random effect. Sulcation Index and sulcal area showed a significant age × diagnosis interaction. **B**. Difference fits ± 95% confidence interval for Sulcation Index and sulcal area. The plots display at approximately what age the mean difference between the predicted fits of the healthy participant and patient groups becomes different from zero and show that the mean difference between the diagnostic groups increases with age.

### Longitudinal allometric analysis: the relationship between annual change in fold metrics and annual change in brain volume in SZ and healthy individuals

The APC of the hull area showed the strongest scaling with the APC of TBV, with no significant differences between the patient and control groups (see Figure 4). In the patient group, the scaling coefficient for the APC of SI and the APC of total brain volume (0.60) was similar to the scaling coefficient for the APC of sulcal area (0.63) and the APC of total brain volume. The patient group showed a significantly stronger scaling, i.e., steeper slope for change in SI and sulcal area compared to the HI group (both p’s <0.001). There were no significant relationships for APCs of sulcal depth and length.

**Figure 4.**
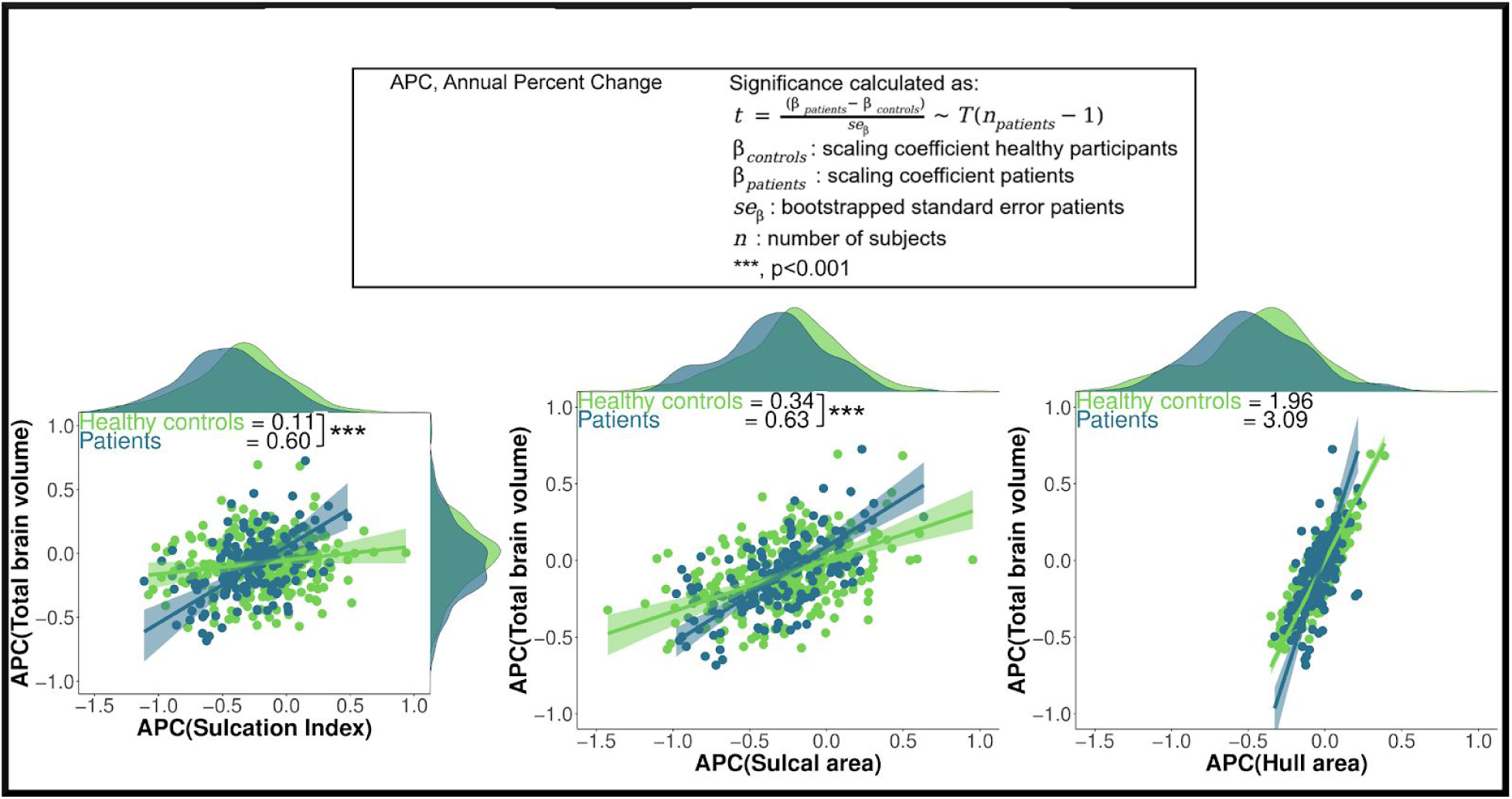
Orthogonal regression fits ± 95% confidence interval and density maps for observed longitudinal allometric scaling of annual percent change (APC) of the folding metrics relative to the APC of total brain volume in the healthy individuals (HI) and the schizophrenia patients in sample 1. The formula for calculating whether the scaling coefficient differed between the HI and patient groups is given. ***=p<0.001 denotes significance after Bonferroni correction for the difference in scaling between the HI group and the patient group. APCs were estimated from values residualized for age, sex, and scanner.

## Discussion

Cross-sectional scaling of the Sulcation Index (SI), sulcal area, and sulcal length with total brain volume (TBV) was significantly higher than expected, i.e. hyperallometric, in healthy individuals (HI), individuals with schizophrenia (SZ) and those with first-episode psychosis (FEP). There were no diagnostic differences in scaling. Longitudinal analyses demonstrated steeper loss of SI and sulcal area through adulthood and beyond in SZ. Finally, we assessed the longitudinal allometric scaling between the annual percent change (APC) of the folding metrics and the APC of TBV. The SZ group showed a significantly stronger scaling between the APC in SI and sulcal area with the APC of TBV compared to the HI group. This finding complemented our previous finding indicating that TBV changes over time are more strongly associated with changes in SI and sulcal area in SZ compared to HI.

### Allometric analyses at baseline: sample 1 and 2

#### Normative scaling coefficients in healthy individuals

The hyperallometric scaling of the SI with TBV was driven by a disproportionate expansion of sulcal area, with increasing TBV accompanied by a proportionate expansion of hull area. Allometric analysis of the sulcal depth and length revealed that disproportionate expansion of sulcal area with increasing TBV arises through a hyperallometric scaling relationship between TBV and sulcal length. The disproportionate hyperallometric scaling between TBV and sulcal length is striking and may be explained by an association between increased TBV and twistier sulci (Germanaud et al., 2012). Whether increased sulcal length in larger brains is also due to the presence of more (‘new’) sulci is difficult to assess. Currently, no common unified classification framework for sulci exists making it challenging to determine whether a sulcus is ‘new’ or not (Ono et al., 1990; Mangin et al., 2010; Mikhael et al., 2018). With respect to the underlying neurobiological mechanisms, it remains unclear what produces the hyperallometric scaling of TBV with sulcal surface area. Possible explanations include an expansion of the progenitor cell pool size producing an increased number of neurons during early-life neurodevelopment and/or as lineage regulation of the different types of cortical progenitors which mediate the tangential dispersion of radially migrating neurons (Rakic, 2009; Ronan et al., 2014; Llinares-Benadero and Borrell, 2019). The robustness of our normative allometric findings are evidenced by the replication in an independent sample and replicating Fish et al. (2017).

#### Comparison of baseline allometric scaling coefficients between healthy individuals and SZ (sample 1) and FEP (sample 2)

Scaling relationships in FEP and SZ are in line with what is expected from the corresponding normative samples. Thus, disproportionate allometric scaling between TBV and SI, sulcal area, and length was also present in individuals with SZ and FEP. In other words, diagnostic differences in global sulcation and its determinants are proportional to the overall effect of diagnosis on TBV, with no added effect of diagnosis per se on these metrics. This suggests that while a diagnosis of SZ or FEP is associated with decreased TBV, global aspects of cortical folding such as SI and its subcomponents are not influenced by the illness. Interestingly, results diverged when using classical methods for controlling for TBV (i.e. normalization and covarying) which do not account for nonlinear relationships between the folding metrics and TBV indicating that accounting for allometry matters when comparing global metrics of cortical folding between HI and individuals with SZ or FEP.

### Longitudinal age trajectories in SZ and healthy individuals (sample 1)

Our longitudinal analyses, to the best of our knowledge, provide the first evidence for diverging lifespan trajectories of SI and sulcal area in SZ when compared to a HI group. After adjusting analyses for the effect of potential non-linear relationships between cortical folding components and TBV on age trajectories, the SZ group demonstrated a steeper SI decline from approximately 32 years onwards and a steeper sulcal area decline from approximately 44 years onwards when compared to the HI group. Our results are in line with increasing *in vivo* evidence for sulcal abnormalities associated with SZ. The amount of cortical surface area depends on cortical curvature. In SZ, sulcal curvature may be disproportionately decreased relative to gyral curvature (Wagstyl et al., 2016). In addition, age-dependent decreases in cortical thickness are more present in sulci than in gyri in HI and this effect may be more pronounced in SZ (Vandekar et al., 2015; Wagstyl et al., 2016). Supragranular layers (layers I-III) are thicker in sulci and thinner in gyri and thus possibly more vulnerable to excessive cortical thinning in SZ (Wagstyl et al., 2016); finally, there is consistent functional as well as post-mortem evidence of supragranular layer pathology in SZ (Harrison and Weinberger, 2005; Williams and Boksa, 2010). Changes in hull surface area may be more sensitive to changes in gyri as compared to sulci and there were no differences in the age trajectories of hull surface area between the SZ and HI groups. Taken together, our results point to abnormal global sulcal changes over time in the SZ group. This may be explained by a disproportionate demyelination in SZ (Mighdoll et al., 2015; Kochunov et al., 2016; Kelly et al., 2018). Tertiary sulci are the latest sulci to develop postnatally and their development is associated with cortical expansion and myelination (Miller et al., 2021). Aging and its associated demyelination may reverse this process, decreasing the prominence of the tertiary sulci leading to a reduction in sulcal area and SI. Nevertheless, future longitudinal studies assessing gyral- and sulcal-specific morphology are necessary to determine whether aberrant morphological changes over time are excessive in sulci.

### Longitudinal allometric analysis: the relationship between annual change in fold metrics and annual change in brain volume in SZ and healthy individuals

This study is the first to show the scaling between annual percent change in subcomponents of SI and change in TBV in HI and individuals with SZ. We do not interpret the reported scaling coefficients from this longitudinal analysis as being hypo-, iso-, or hyperallometric as there are, to the best of our knowledge, no prior studies presenting normative longitudinal scaling coefficients for folding subcomponents have been published. Scaling coefficients for SI and sulcal area were higher in the SZ group compared to the HI group, i.e. TBV changes over time are more strongly associated with changes in SI and sulcal area in SZ compared to HI. This complements our findings of the group differences in age trajectories. Loss of SI and sulcal area over time was higher in SZ and this loss has an atypical strong association with change in TBV. As such it may be that loss of SI and sulcal area, rather than hull area, sulcal depth and length, are important contributors to the disproportionate loss of TBV over time in SZ.

A number of limitations should be taken into account when interpreting these findings. In sample 1 antipsychotics usage information was heterogeneous and unfortunately limited. Thus, the obtained correlations between measures of chlorpromazine equivalents and brain metrics reported in this study are approximate and the results must be interpreted cautiously. Recent work demonstrated a power law relation between average global cortical thickness, total exposed cortical surface area and total cortical surface area which describes the degree of folding (Wang et al., 2016). Whether this power law also predicts the relationship between the metrics used in the current study should be addressed in future studies. We assessed global but not regional (e.g. at the level of the individual sulcus) folding. By examining global aspects of cortical folding we address brain-wide geometrical organization and as such provide a building block for future studies examining more local characteristics of folding.

## Supplement

### Image quality assurance

#### Sample 1

After the completion of the preprocessing pipeline for all T1 MRI scans in FreeSurfer, we rigorously assessed the quality of the data using a combination of visual inspection and the examination of several quantitative metrics following recent recommendations from the literature (Rosen et al., 2018). First, we calculated five quality measurements based on the proposed by the Preprocessed Connectome Project (http://preprocessed-connectomes-project.org/quality-assessment-protocol/) to identify images that were unusable: signal-to-noise ratio (SNR), contrast to noise ratio (CNR), foreground to background Energy Ratio (FBER) and percent artefact Voxels (Artifacts) and Entropy Focus Criterion (Entropy). As in (Vandekar et al., 2015) we defined the threshold for outliers as [mean-(2.698*SD)] for SNR, CNR and FBER metrics; and Artifacts and Entropy were only tested for [mean-(2.698*SD)] based on the exclusion criteria explained by ENIGMÁs QA protocol. Next, for each scan the whole brain mean cortical thickness, total surface area, total white matter volume, total grey matter volume, subcortical grey matter volume and intracranial volume were calculated. Then, we summed for each FreeSurfer directory the amount of ENIGMA outliers over all measures and calculated the mean+(2.698*SD) for the amount of outliers over the whole sample and designate those above the threshold as outliers. These quality control steps resulted in the below exclusions.

Following the exclusion of participants that met our exclusion criteria (585 scans) a sample of 1239 scans was obtained. After the evaluation of the computed quantitative measures of image quality, 152 scans were considered of insufficient quality and were excluded from our analysis (see SFigure 1A). These excluded scans were validated by manually assessing image quality after preprocessing to assure that all preprocessing steps worked and to avoid the propagation of error along the analysis. The parameters that were most useful for objectively detect artefacts visually were agreed on between researchers, e.g. incorrect sulcal labelling or insufficient quality of sulcal segmentation resulting in gross anatomical abnormalities. These visual checks were performed in BrainVisa. Those scans with poor initial sulcal labelling were manually edited, which often resulted in better sulcal labelling when manually inspected (see SFigure 1B for examples of “good”, “medium” and “bad” quality sulcal labelling after first preprocessing of the images). In addition, 14 scans were excluded after this manual inspection.

#### Sample 2

QA information for sample 2 can be found in (Pina-Camacho et al., 2016). Briefly, manual edits, mainly for errors in skull-stripping, were made. Thereafter, no outliers for FreeSurfer processed images were detected. However, manual inspection of the BrainVisa (see above) output led to the exclusion of 32 participants.

### Sulcation Index and Gyrification Index

The sulcation index (*SI*) is the ratio between total sulcal surface area and hull area:

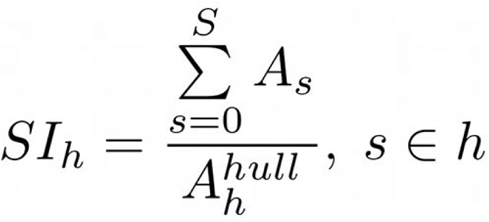

where *h* yields for hemisphere, *s* for sulcus, and *A* is area, being 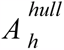 the area of the brain hull for hemisphere *h*. The SI differs from the gyrification index (GI) (Zilles et al., 1988)which is calculated as:

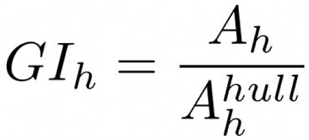

where *h* yields for hemisphere, and *A* is area, being 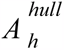 the area of the cortex for hemisphere *h*. We made a diagram which gives additional information about the differences between SI and GI:

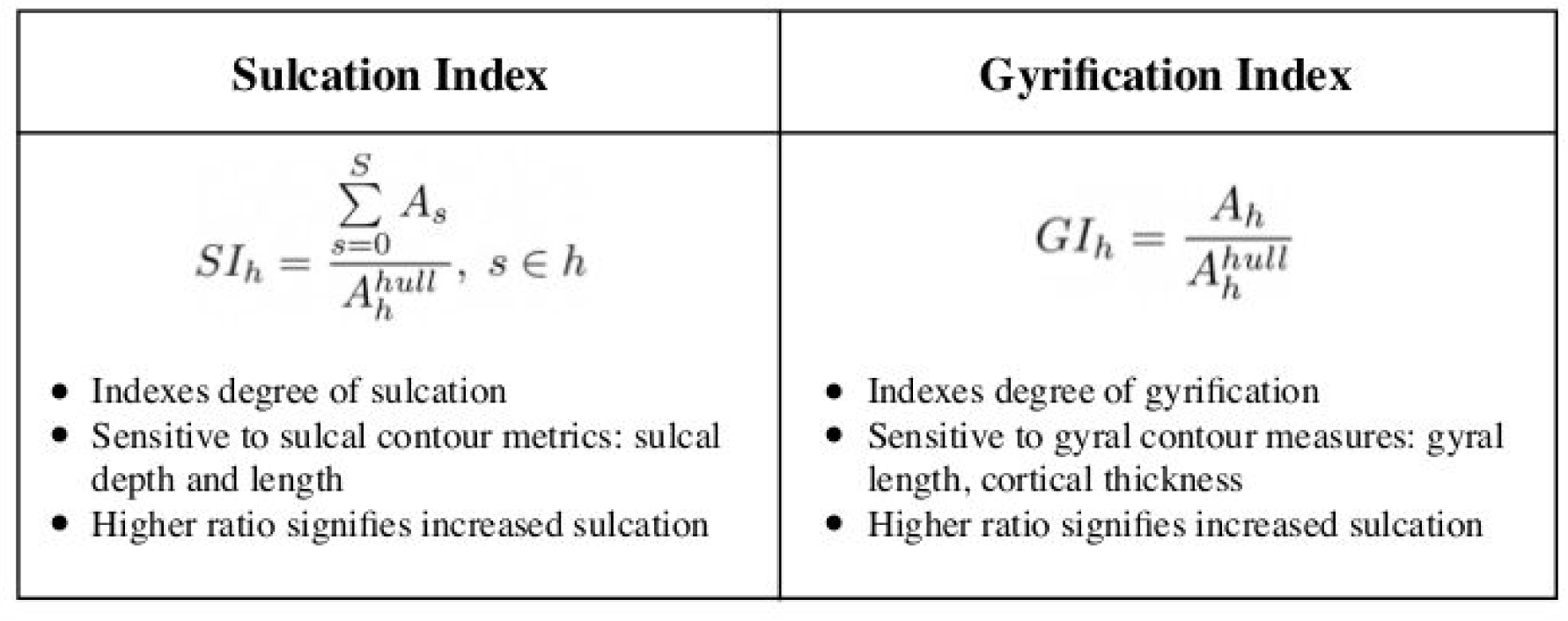

| Sulcation Index | Gyrification Index |
| --- | --- |
| $SI_h = \frac{\sum_{s=0}^S A_s}{A_h^{hull}}, s \in h$<br>• Indexes degree of sulcation<br>• Sensitive to sulcal contour metrics: sulcal depth and length<br>• Higher ratio signifies increased sulcation | $GI_h = \frac{A_h}{A_h^{hull}}$<br>• Indexes degree of gyrification<br>• Sensitive to gyral contour measures: gyral length, cortical thickness<br>• Higher ratio signifies increased sulcation |

The SI’s particular sensitivity to global changes in sulcal morphology as opposed to changes in gyri was the reason for assessing SI. A cortex with extensive folding has a large *SI*, whereas a cortex with low degree of folding has a small *SI*. At a constant outer cortex area, the *SI* increases with the number and area of sulcal folds, whereas the *SI* of a lissencephalic cortex is zero.

---

**SFigure 1.**
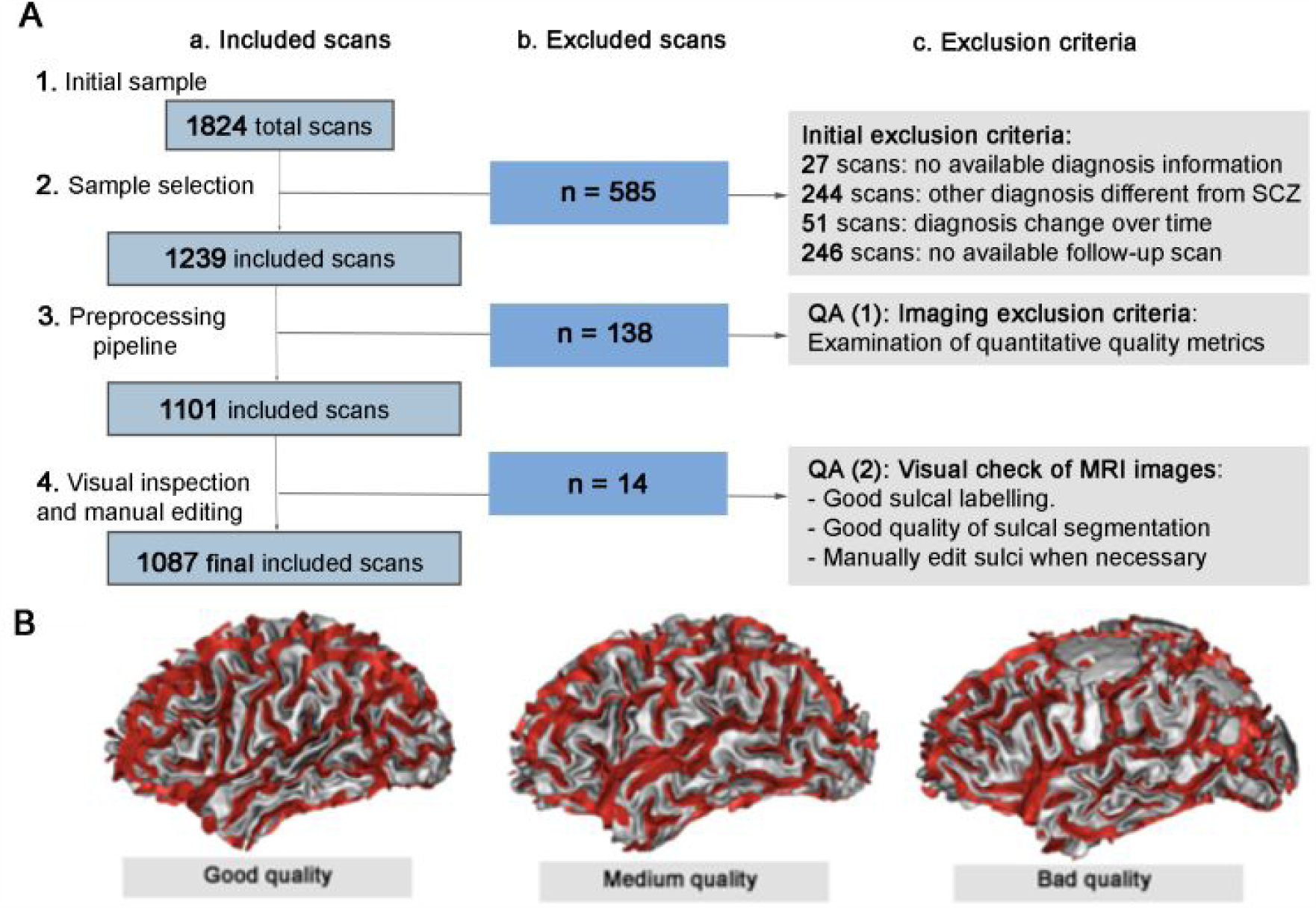
**A**) Flowchart summarizing the exclusion criteria process: from an initial sample of 1824 total scans, a total of 737 scans were excluded for subsequent analysis. The final sample comprises 1087 scans: a. indicates the number of included scans; b. the number of excluded scans and c. the exclusion criteria (initial and imaging criteria) applied to the sample. **B**) Illustrates examples of sulcal segmentation of different qualities performed with Brainvisa: good quality of sulcal reconstruction, medium quality of sulcal reconstruction, included after manual sulcal editing; and bad quality scan which shows a lump caused by incorrect sulcal reconstruction which caused exclusion.

**SFigure 2.**
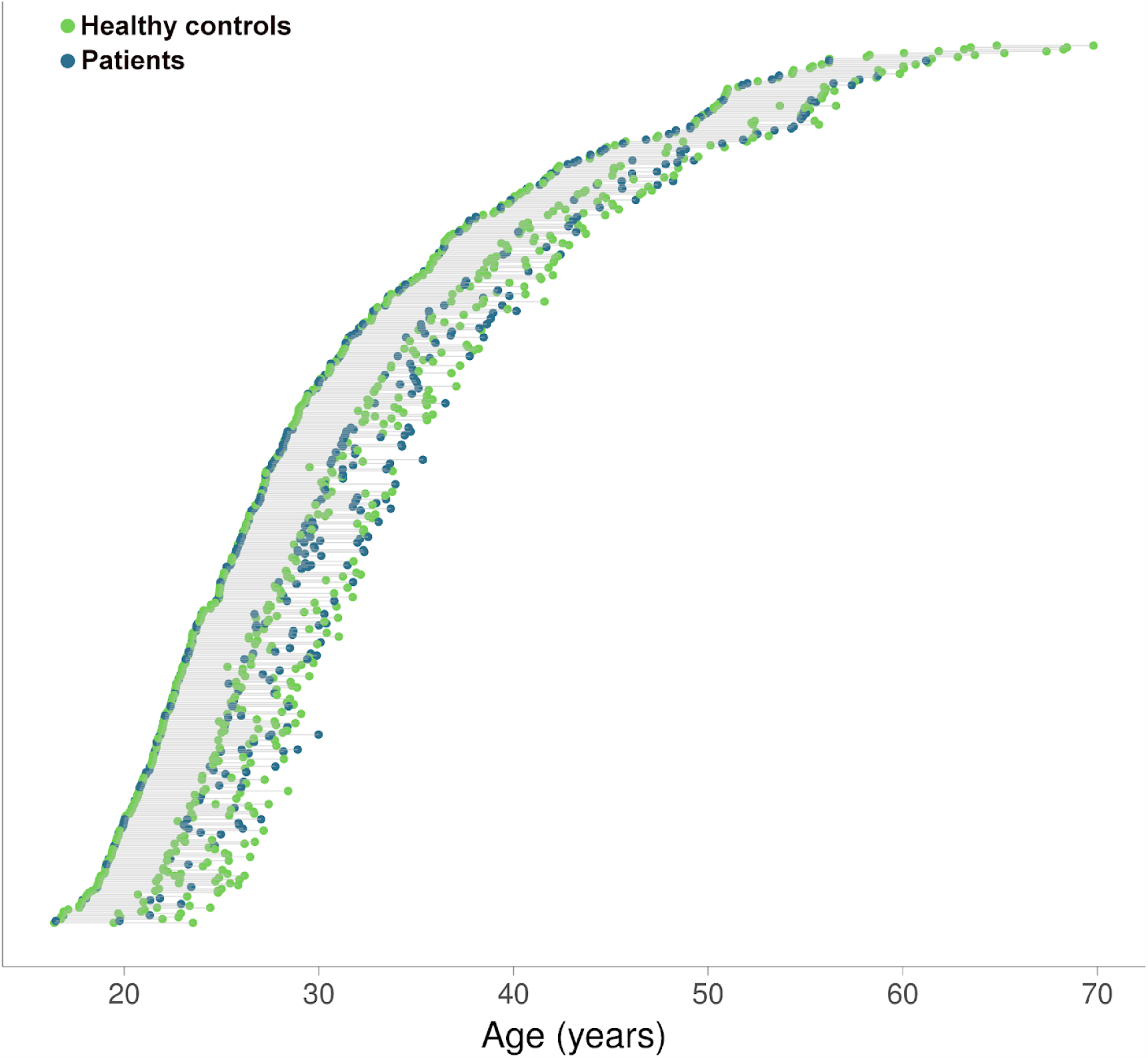
Age at scans for all participants from sample 1. Each of the 1087 scans obtained is represented by a dot; each of the 466 participants is shown in a different row, with their two or three scans connected by a straight grey line. Healthy individuals and patients are marked separately.

**SFigure 3.**
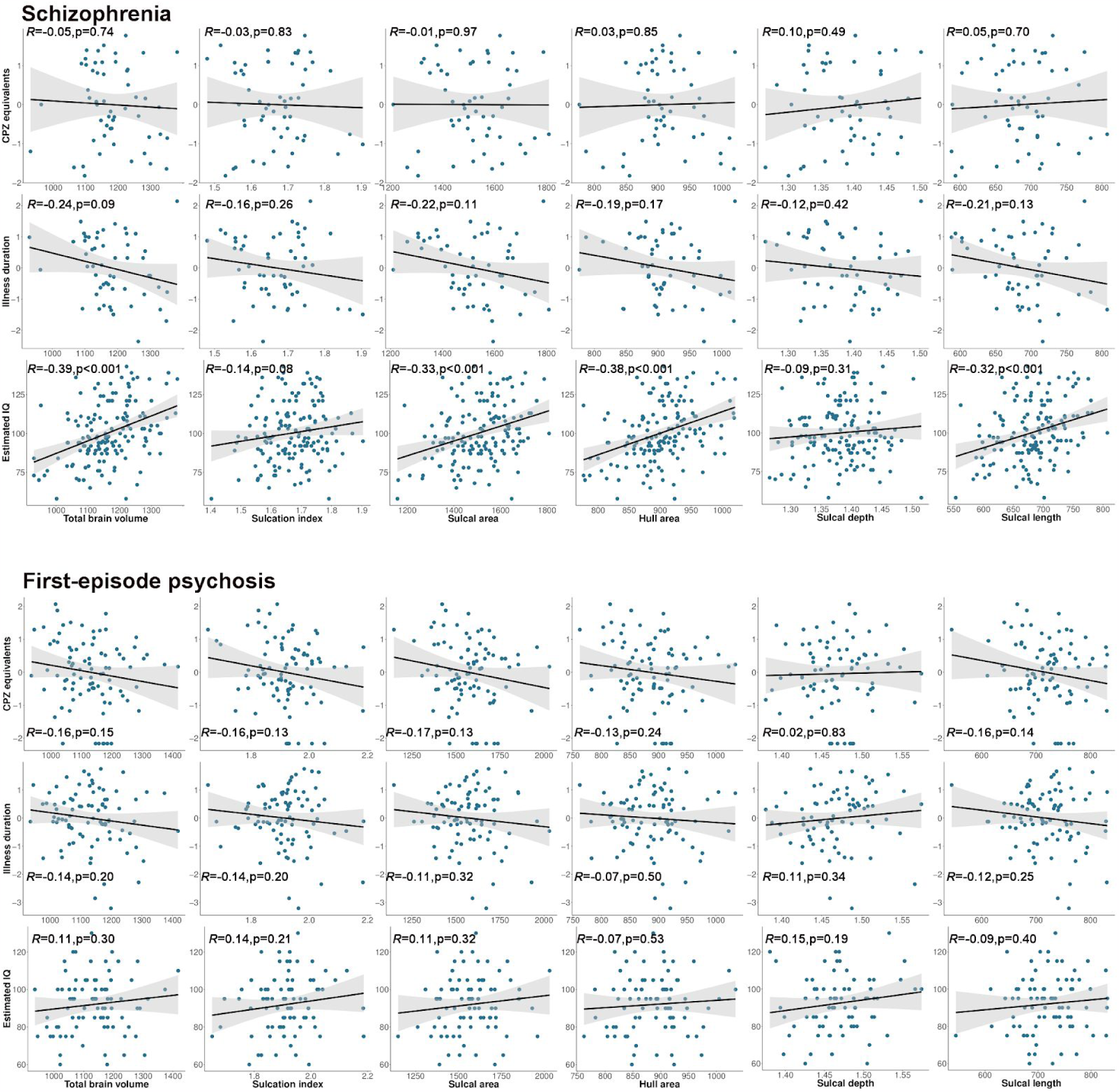
Regression plots for patients from sample 1 and 2 showing the relationship between the brain measures at baseline and cumulative antipsychotics intake, illness duration, and estimated IQ. Chlorpromazine equivalents and illness duration were first normalized by selecting the optimal transformation method, based on the lowest Pearson p test statistic for normality, among the Yeo-Johnson, exp(x), log10, square-root, arcsinh and orderNorm transformations (bestNormalize package, R). Participants missing data in sample 1: cumulative antipsychotic usage (103), illness duration (60) and estimated IQ (44). Pearson correlations for each pair of variables were not significant after Bonferroni correction across samples. CPZ, chlorpromazine equivalents; IQ, intelligence quotient. CPZ, chlorpromazine; IQ, intelligence quotient.

